# Shared immune signatures in acute myocardial infarction and periodontitis: the role of FCN1 and LYN as potential biomarkers

**DOI:** 10.64898/2026.07.23.740187

**Authors:** Kun Zhang, Yanqing Wang

**Author notes:** Corresponding Author: Yanqing Wang.

## Abstract

**Background:** Acute myocardial infarction (AMI) and periodontitis (PD) have been epidemiologically linked, but the molecular mechanisms underlying this association remain elusive. We aimed to elucidate shared pathogenic signatures between AMI and PD using a comprehensive bioinformatics approach.

**Methods:** We integrated transcriptomic data from multiple Gene Expression Omnibus datasets, including an AMI cohort profiled from enriched circulating endothelial cells (GSE66360) with whole-blood validation (GSE48060), and PD gingival tissue cohorts (GSE16134 and GSE10334). Weighted gene co-expression network analysis, protein-protein interaction network analysis, and LASSO-based feature selection were applied. Functional enrichment, immune deconvolution, transcription factor-gene regulatory network analysis, and single-cell RNA sequencing analysis were performed to characterize shared molecular features.

**Results:** We identified 95 shared differentially expressed genes (DEGs) between AMI and PD. By intersecting the shared upregulated DEGs with disease-associated WGCNA modules, we obtained 46 candidate shared genes. LASSO-based feature selection further highlighted FCN1 and LYN as overlapping candidates, which showed good discriminative performance in both training and external validation cohorts in ROC analyses. Enrichment analyses suggested that the shared signature was mainly related to myeloid cell migration, phagocytosis, and neutrophil-related inflammatory pathways (e.g., neutrophil extracellular trap formation). Immune deconvolution in PD gingival tissues suggested increased plasma cells and neutrophils and decreased resting memory CD4+ T cells; immune deconvolution results in the AMI cohort were interpreted cautiously due to the CEC-enriched sample source. Single-cell analysis revealed that FCN1 and LYN were predominantly expressed in macrophage/monocyte-derived populations.

**Conclusions:** Our study suggests a shared inflammatory and immune-mediated transcriptomic program linking AMI and PD, and identifies FCN1 and LYN as candidate shared immune markers. These findings provide a molecular rationale for the oral-cardiovascular association and are hypothesis-generating, warranting future experimental and prospective validation.

## Introduction

Cardiovascular diseases (CVDs) remain the leading cause of morbidity and mortality worldwide, with acute myocardial infarction (AMI) representing a particularly severe manifestation ^1^. Characterized by severe retrosternal pain and potentially life-threatening complications such as cardiac arrhythmia and heart failure, AMI poses a significant global health burden ^2^. Despite advances in prevention and treatment, the incidence of AMI continues to rise, particularly among middle-aged and elderly individuals, as well as high-risk groups including those with pre-existing heart conditions, smokers, diabetics, and individuals with hyperlipidemia ^3^.

Concurrently, periodontitis (PD), a chronic inflammatory disease primarily caused by the destruction of periodontal tissue by oral bacteria, has emerged as a significant public health concern ^4^. Recent evidence suggests that PD’s impact extends beyond oral health, potentially contributing to various systemic illnesses through mechanisms involving pro-inflammatory cytokines, bacterial dissemination, and immune system modulation ^5,6^. However, the systemic implications of PD remain underappreciated in many healthcare systems globally.

The potential link between oral health and cardiovascular disease has garnered increasing attention in recent years. Epidemiological studies have consistently demonstrated an association between PD and CVDs, particularly AMI ^7,8^. A landmark study utilizing the functional periodontal pentagon risk diagram (PPRD) revealed a significant correlation between PPRD scores and AMI status, with radiographic evidence of bone loss emerging as the strongest individual predictor ^9^. More recently, a comprehensive Korean nationwide cohort study provided compelling evidence for a causal relationship between severe periodontitis (SPD) and increased risk of AMI, stroke, and major adverse cardiovascular events (MACE) ^10^. These findings were further corroborated by a case-control study demonstrating a dose-dependent association between PD severity and AMI risk ^11^.

Despite the accumulating epidemiological evidence, the precise molecular mechanisms underlying the PD-AMI association remain elusive. The advent of high-throughput genomic technologies and advanced bioinformatics approaches offers unprecedented opportunities to explore the shared pathogenesis of these conditions at a molecular level. Identifying common transcriptional patterns and regulatory networks between AMI and PD could not only elucidate the biological basis of their association but also reveal novel therapeutic targets and diagnostic biomarkers. The complex, multifactorial nature of both AMI and PD necessitates an integrative, systems-level approach to unravel their shared pathogenic mechanisms. Recent studies have highlighted the role of chronic inflammation, immune dysregulation, and alterations in lipid metabolism as potential common pathways ^12,13^. However, a comprehensive, multi-omics analysis comparing the molecular landscapes of AMI and PD is lacking.

Although AMI and PD have been consistently linked epidemiologically, reusable shared immune transcriptomic signatures that connect these conditions remain poorly defined. Here, we integrated multiple transcriptomic datasets of AMI and PD (gingival tissues) using differential expression analysis, WGCNA, PPI network analysis, and LASSO-based feature selection to delineate shared inflammatory pathways. The flowchart of the study was detailed in Figure 1(Fig. 1).

**Fig. 1.**
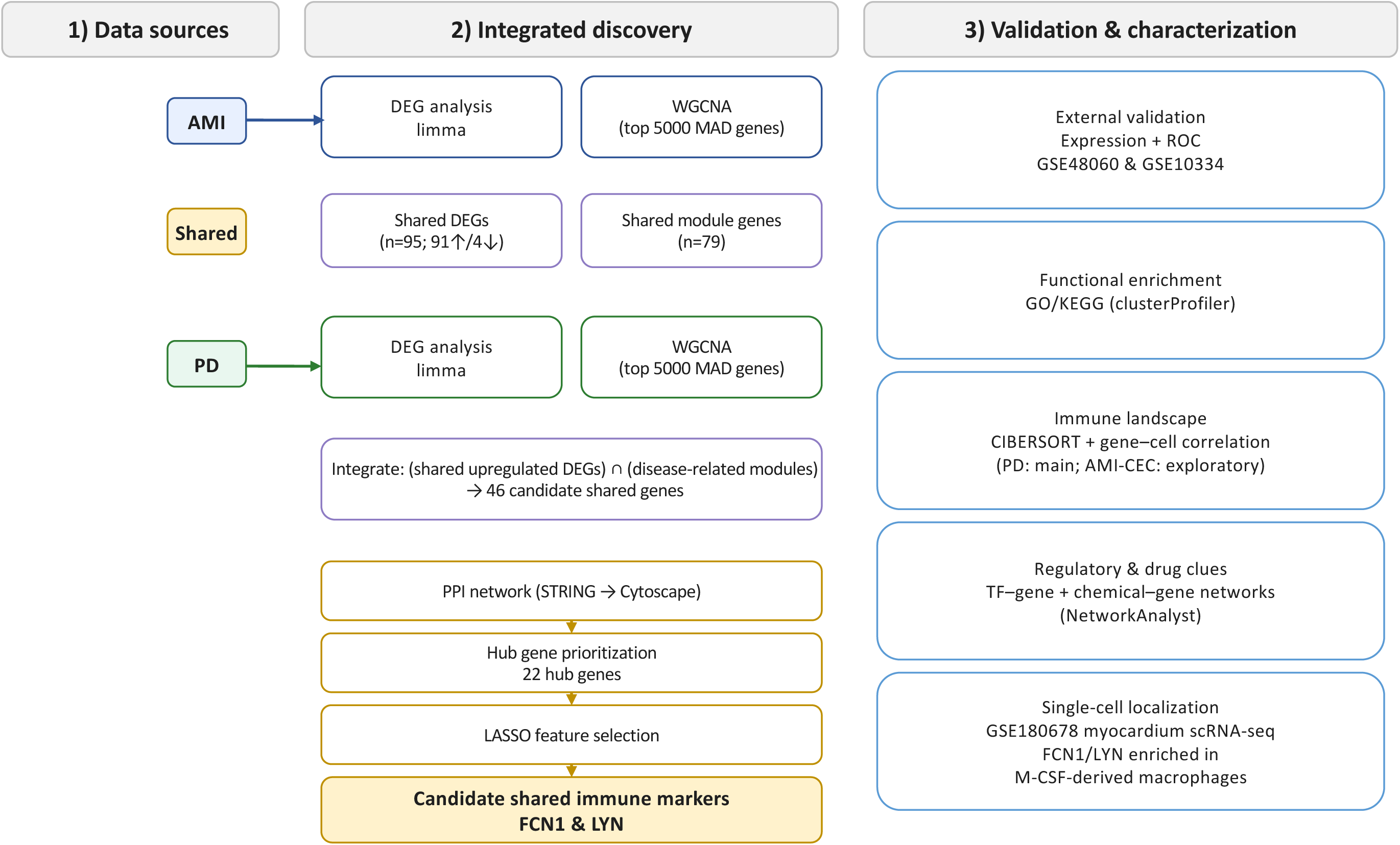
Workflow diagram for the whole study.

## Materials and Methods

### Data Preparation

For AMI, GSE66360 (enriched circulating endothelial cells isolated from peripheral blood; 49 AMI and 50 controls) was used as the training cohort, and GSE48060 (peripheral blood; 31 AMI and 21 controls) served as an external validation cohort. For PD, GSE16134 (gingival tissues; 241 PD and 69 controls) was used for training and GSE10334 (gingival tissues; 183 PD and 64 controls) for validation. In addition, the single-cell dataset GSE180678 (AMI myocardium) was used to localize candidate markers at the cellular level ^14^. Detailed dataset information is provided in Table 1.

**Table 1.** Datasets information.

| GSE series | Disease | Samples | Source types | Platform |
| --- | --- | --- | --- | --- |
| GSE66360 | AMI | 49 AMI and 50 controls | Whole Blood | GPL570 |
| GSE48060 | AMI | 31 AMI and 21 controls | Blood | GPL570 |
| GSE16134 | PD | 241 PD and 69 controls | Gingival tissues | GPL570 |
| GSE10334 | PD | 183 PD and 64 controls | Gingival tissues | GPL570 |
| GSE180678 | AMI | 1 AMI | Myocardium (single cell RNA sequence) | GPL20301 |

### Identification of Differentially Expressed Genes

To identify differentially expressed genes (DEGs) in the AMI and PD datasets (GSE66360 and GSE16134), we employed the “Limma” package in R (version 4.4.1) ^15^. Genes were considered differentially expressed if they met two criteria: an absolute log2 fold change greater than 0.585 and an adjusted p-value less than 0.05 (|log2 FC| > 0.585 and adj.p-value <0.05). We visualized the DEG results using volcano plots created with the “ggplot2” package in R ^16^. To identify common DEGs between AMI and PD, we used the “VennDiagram” package ^17^ to determine the intersection of the DEG sets from both analyses.

### Weighted gene co-expression network analysis (WGCNA)

We performed WGCNA to identify clusters of highly correlated genes and their associations with disease states. Using the “Seurat” package ^18^, we selected the top 5000 highly variable genes from each dataset for analysis. The WGCNA was conducted using the “WGCNA” package ^19^, following these steps: determination of soft thresholds using the pickSoftThreshold function, construction of weighted adjacency matrices, and calculation of correlations between gene modules and disease states. The module showing the highest correlation with grouped traits was identified as the key module. We then used the “VennDiagram” package to identify the intersection between WGCNA results and previously identified DEGs, defining these as candidate genes for further analysis.

### Protein-protein interaction (PPI) Network Analysis

To explore potential interactions among the proteins encoded by our candidate genes, we utilized the Search Tool for the Retrieval of Interacting Genes (STRING) database (Version 12.0, http://string-db.org). We set a combined score > 0.4 as the threshold for medium confidence interactions. The resulting PPI network was visualized and analyzed using Cytoscape software (version 3.10.2) ^20^. From this network, we identified the twenty-two most significant hub genes based on their connectivity and centrality measures.

### Functional Enrichment Analysis

To understand the biological context of our identified hub genes, we performed Gene Ontology (GO) and Kyoto Encyclopedia of Genes and Genomes (KEGG) pathway enrichment analyses in R using the clusterProfiler package. Gene identifiers were converted when necessary (e.g., with org.Hs.eg.db). GO analysis included evaluations of biological processes (BP), cellular components (CC), and molecular functions (MF) ^21^. Enrichment results with p < 0.05 were considered statistically significant and visualized using ggplot2 (and related plotting functions).

### Machine learning-assisted identification of candidate diagnostic markers

We applied the Least Absolute Shrinkage and Selection Operator (LASSO) regression as a feature selection approach to prioritize candidate diagnostic markers from the 22 hub genes identified in our previous analyses ^22^. Genes with non-zero coefficients were retained as candidate markers. Discriminative performance was evaluated by Receiver Operating Characteristic (ROC) curves and the area under the curve (AUC) in the training cohorts, and robustness was further assessed in two independent validation datasets (GSE48060 for AMI and GSE10334 for PD).

### Immune infiltration analysis

We estimated immune cell composition in case and control groups using the CIBERSORT algorithm ^23^, which infers the relative proportions of immune cell types from bulk gene expression data. We then assessed the correlation between candidate marker expression and the inferred immune cell fractions using Spearman’s rank correlation coefficient. Results with p < 0.05 were considered statistically significant and visualized using ggplot2. Because the PD datasets represent bulk gingival tissues, immune deconvolution results in PD were interpreted directly. In contrast, GSE66360 profiles enriched circulating endothelial cells; therefore, immune deconvolution in this AMI cohort was treated as exploratory and interpreted cautiously.

### Transcription Factor and Chemical Interaction Analysis

To identify potential regulatory mechanisms, we predicted transcription factors (TFs) and chemicals interacting with our key genes using NetworkAnalyst 3.0 (https://www.networkanalyst.ca) ^24^. This analysis helps elucidate the upstream regulators and potential drug interactions of our identified genes, providing insights into their broader biological context and potential therapeutic implications.

### Single-cell sequencing analysis

We conducted single-cell RNA sequencing analysis using the Seurat package in R, following established protocols. Our analysis procedures included rigorous quality control, retaining cells with 500-4000 unique genes and less than 15% mitochondrial gene expression. We normalized the count matrix to account for technical variations, identified the top 2000 highly variable genes using variance-stabilizing transformation, and integrated data across samples to mitigate batch effects. Dimensionality reduction was performed using Principal Component Analysis, followed by cell clustering to identify distinct populations. We visualized cellular heterogeneity using t-distributed Stochastic Neighbor Embedding (t-SNE) and annotated cell clusters using the celldex package. Throughout the analysis, we used default parameters unless otherwise specified, with all computational work performed in R. This comprehensive approach enabled us to characterize the cellular composition and gene expression patterns at single-cell resolution, providing valuable insights into the underlying biology of our samples.

## Results

### DEGs in patients with AMI and PD

AMI datasets contained circulating endothelial cells (CECs) isolated from peripheral blood, comprising 99 samples (50 controls, 49 AMI). With criteria of adjusted p-value < 0.05 and |logFC| > 0.585, we identified 1176 differentially expressed genes (DEGs), including 681 upregulated and 495 downregulated genes (Fig. 2A). For PD, we analyzed the GSE16134 dataset, which included 310 gingival tissue samples (69 controls, 241 PD). Applying identical criteria, we identified 1038 DEGs in PD, with 687 upregulated and 351 downregulated genes (Fig. 2B). To identify common genetic signatures between AMI and PD, we intersected the DEGs from both datasets, revealing 95 shared DEGs. Among these shared DEGs, 91 were consistently upregulated in both conditions (Fig. 2C), while 4 were consistently downregulated (Fig. 2D).

**Fig. 2.**
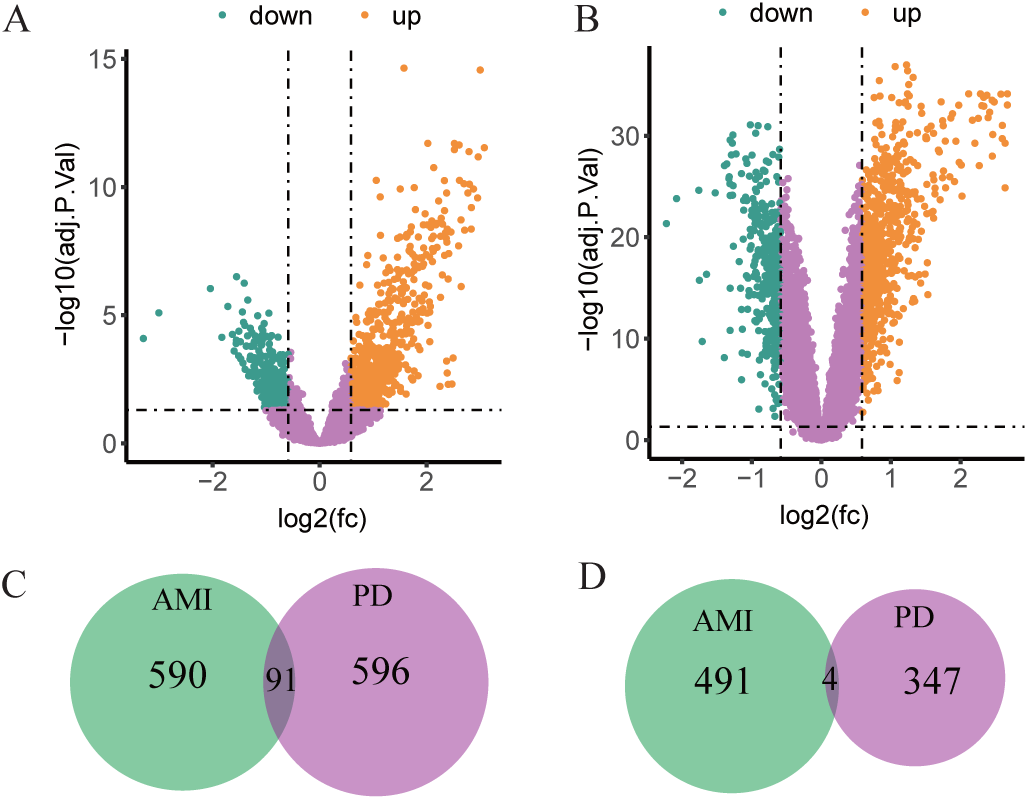
Identification of differentially expressed genes (DEGs). (A) DEGs between AMI vs. control groups. (B) DEGs between PD vs. control groups. Downregulated and up-regulated DEGs are shown in green dots and orange dots correspondingly. Gray dots represent no statistically significant genes. The cut-off criterion is adj.pvalue < 0.05 and |log2FC|> 0.585. (C-D) Venn plot identifies co-upregulated and co-downregulated DEGs.

### WGCNA network construction and module identification

In WGCNA, we retained the top 5000 genes ranked by median absolute deviation (MAD) in each dataset to focus on informative variation. We selected soft thresholds of β = 7 for AMI and β = 12 for PD (scale-free R2 = 0.9) to ensure scale-free networks (Fig. 3A-D). This analysis revealed 8 modules in the AMI dataset (GSE66360, Fig. 3E) and 13 modules in the PD dataset (GSE16134, Fig. 3F). We then assessed the correlation between these modules and disease states using Pearson correlation coefficients. In the AMI dataset, the yellow module showed the strongest positive correlation with disease (r = 0.66), while the black module exhibited the most significant negative correlation (r = −0.29; Fig. 3E). For PD, the purple module demonstrated the highest positive association (r = 0.67), and the mediumpurple3 module showed the strongest negative correlation (r = −0.61; Fig. 3F). Interestingly, we found 79 overlapping genes between the most positively correlated modules of both diseases, but no overlap in the negatively correlated modules (Fig. 3G). Further analysis using Venn diagrams revealed 46 candidate genes that were common between the WGCNA results and the upregulated DEGs in both AMI and PD (Fig. 3H). These findings suggest potential shared molecular mechanisms between AMI and PD.

**Fig. 3.**
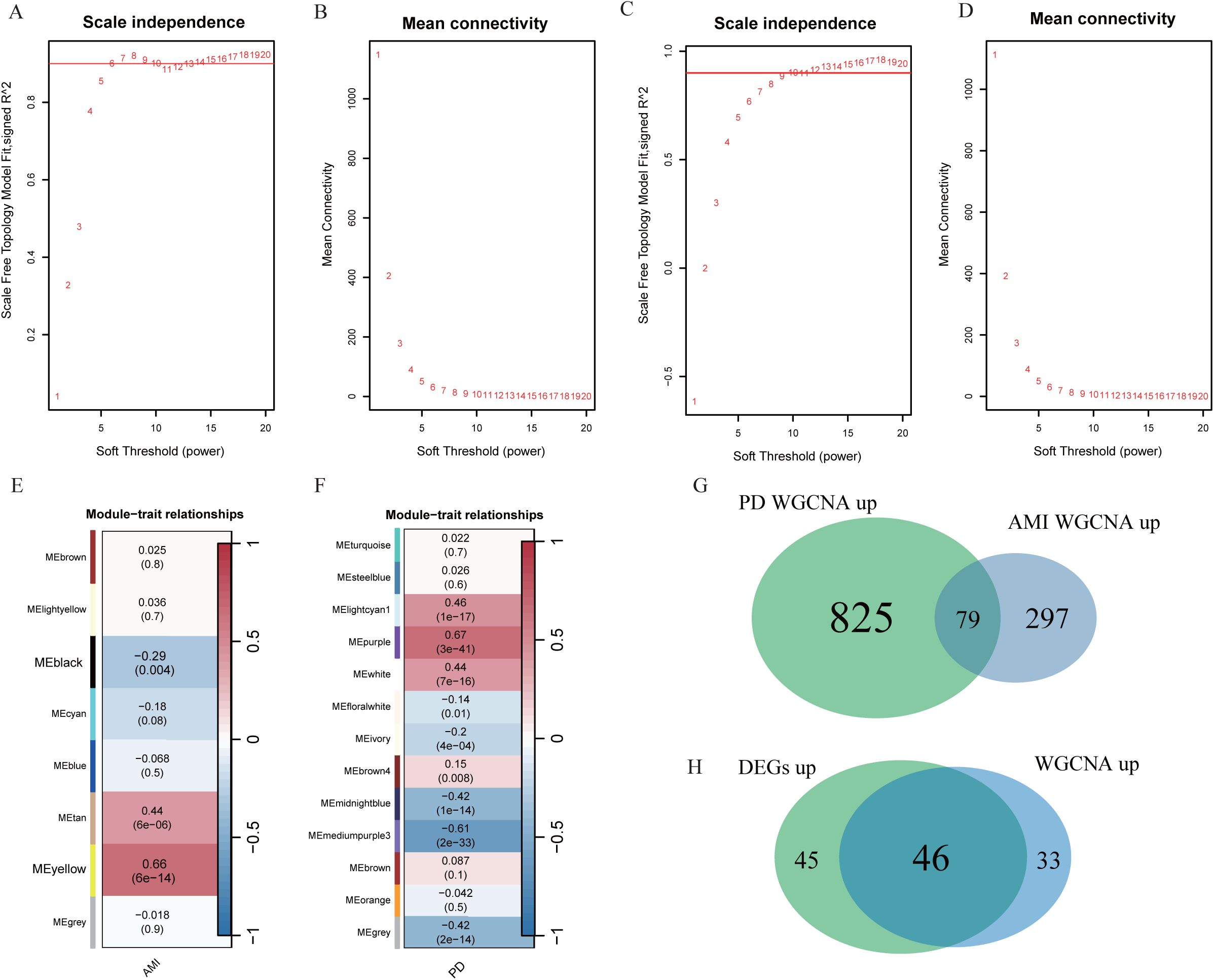
WGCNA analysis. (A-D) Mean connectivity for scale independence and soft threshold in the AMI and the PD datasets. (E-F) Heatmap of the correlation analysis of module eigengenes with clinical phenotypes in AMI and PD. Red color represents positive correlation and blue color represents negative correlation. (G) Venn plot identifies intersection genes between yellow module in AMI and purple module in PD. (H) Venn plot identifies intersection genes between co-upregulated DEGs and the intersection in (G)

### PPI and hub genes’ identification

We utilized PPI network to identify key regulatory genes. After excluding non-protein-coding genes, we used 41 common genes to build a PPI network using the STRING database. The resulting network comprised 41 nodes interconnected by 264 PPI edges (Fig. 4). We then imported this network into Cytoscape for further analysis. Employing the Degree algorithm, we identified the top 22 genes with a Degree score greater than 28. These genes emerged as central nodes in the network.

**Fig. 4.**
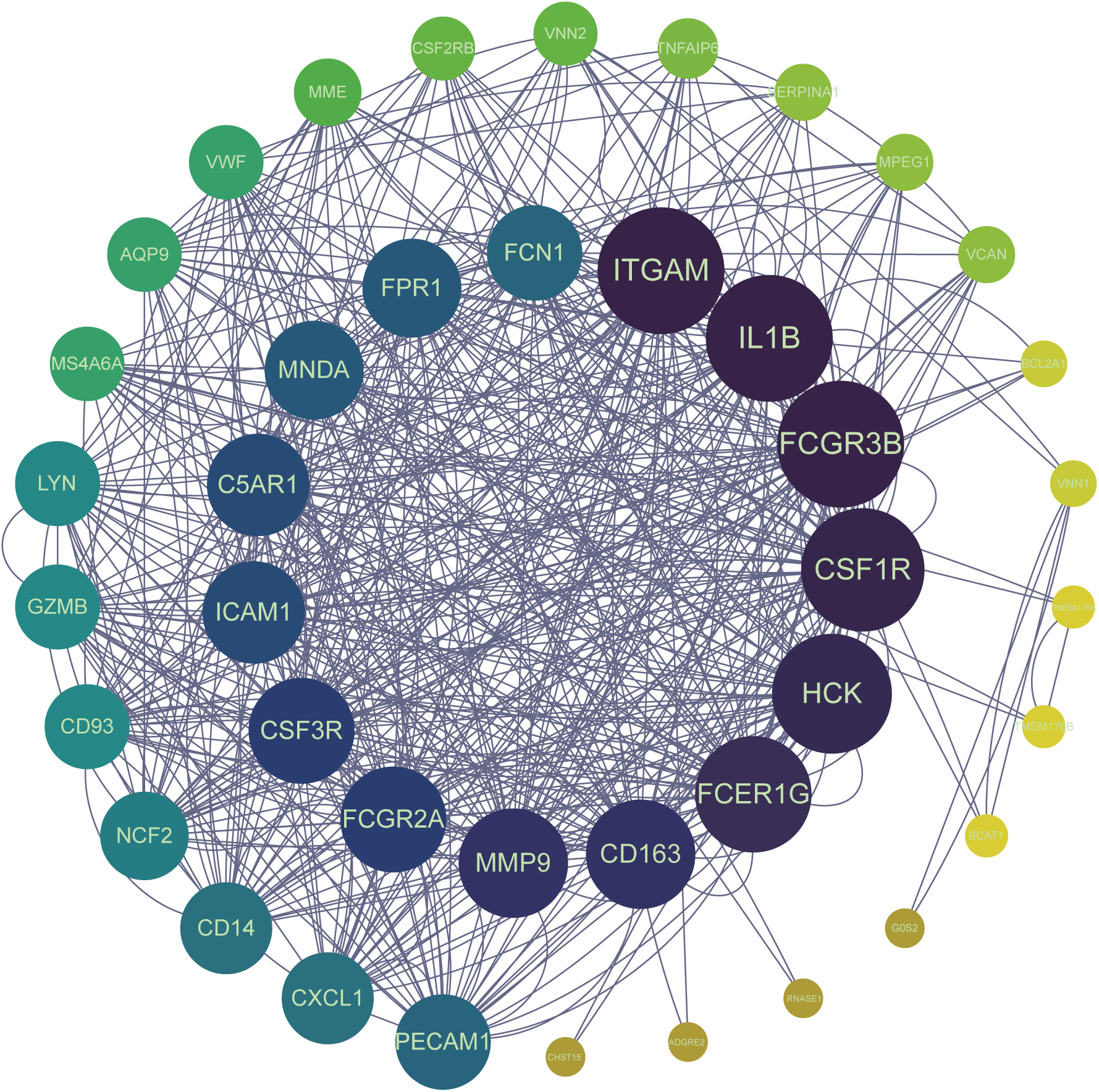
The PPI network of 41 common genes between AMI and PD. The circular node represents genes and the edge represents the interaction between nodes.

### Enrichment analysis

To elucidate the functional roles and biological significance of the 22 identified hub genes, we performed GO and KEGG pathway analyses. The GO analysis, encompassing BP, CC, and MF, revealed several significantly enriched terms. In the BP category, “leukocyte migration,” “phagocytosis,” “myeloid leukocyte migration,” and “activation of immune response” were prominently featured. The MF analysis highlighted “immune receptor activity,” “IgG binding,” and “immunoglobulin binding” while the CC analysis identified “secretory granule membrane,” “external side of plasma membrane,” “tertiary granule,” “ficolin-1-rich granule” as key cellular locations for these genes’ products (Fig. 5). Complementing these findings, our KEGG pathway analysis uncovered significant enrichment in “Staphylococcus aureus infection,” “Tuberculosis,” and “Neutrophil extracellular trap formation” pathways. These results collectively suggest that the shared molecular mechanisms between AMI and PD are predominantly related to immune response regulation, cellular adhesion, and inflammatory processes, particularly those involving bacterial infections.

**Fig. 5.**
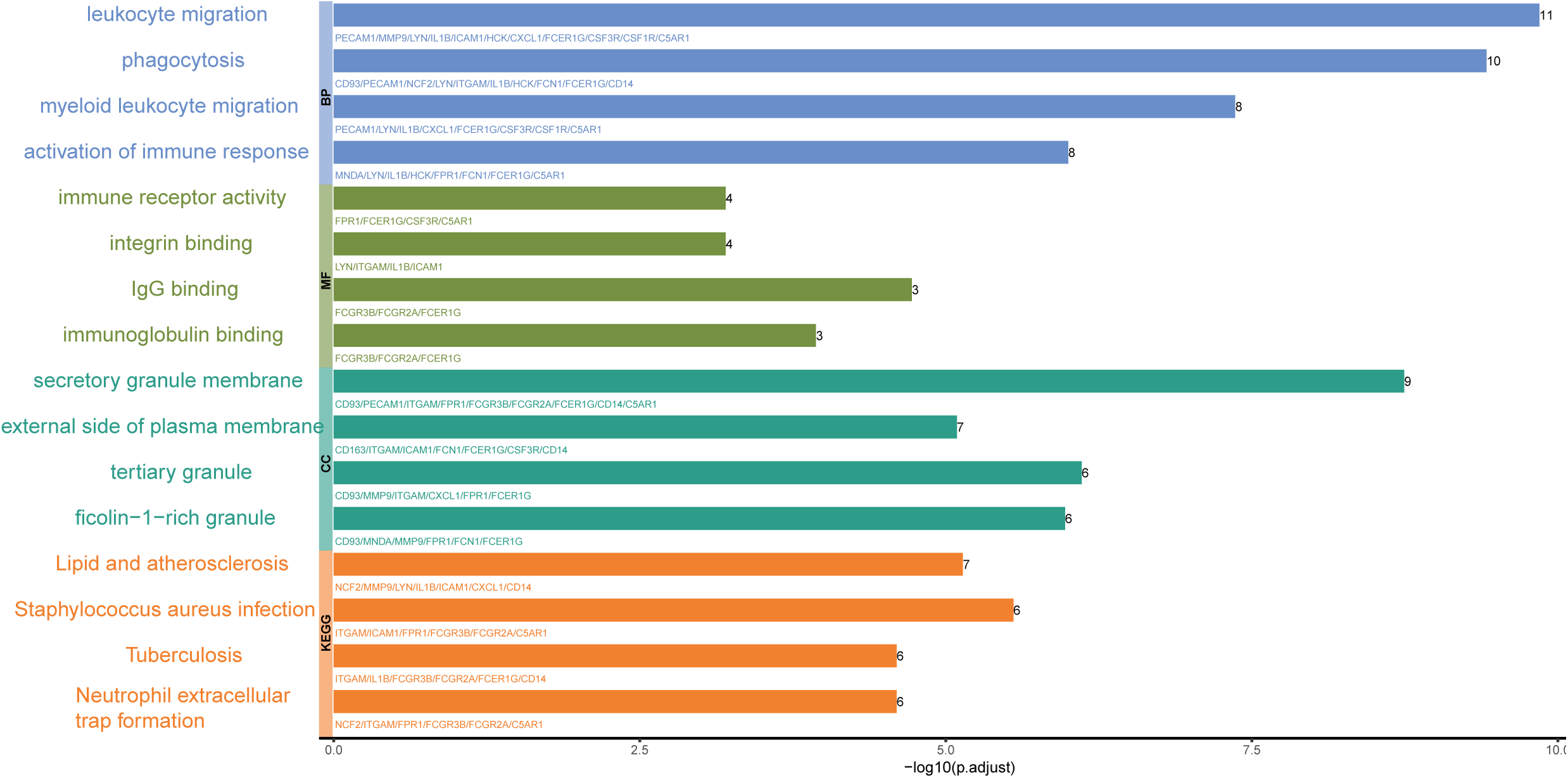
GO and KEGG enrichment analysis of hub genes.

### Identification of potential shared diagnostic genes

To prioritize candidate shared diagnostic markers, we applied LASSO regression to the 22 hub genes separately in the AMI (GSE66360) and PD (GSE16134) training cohorts (Fig. 6). This analysis revealed ten candidate markers in the AMI cohort (IL1B, HCK, FCER1G, CD163, MMP9, ICAM1, MNDA, FCN1, LYN, GZMB) and seven candidate markers in the PD cohort (CSF3R, FPR1, PECAM1, FCN1, CXCL1, LYN, CD93). FCN1 and LYN were selected in both models, suggesting that they may represent shared immune-related candidate markers across the two conditions.

**Fig. 6.**
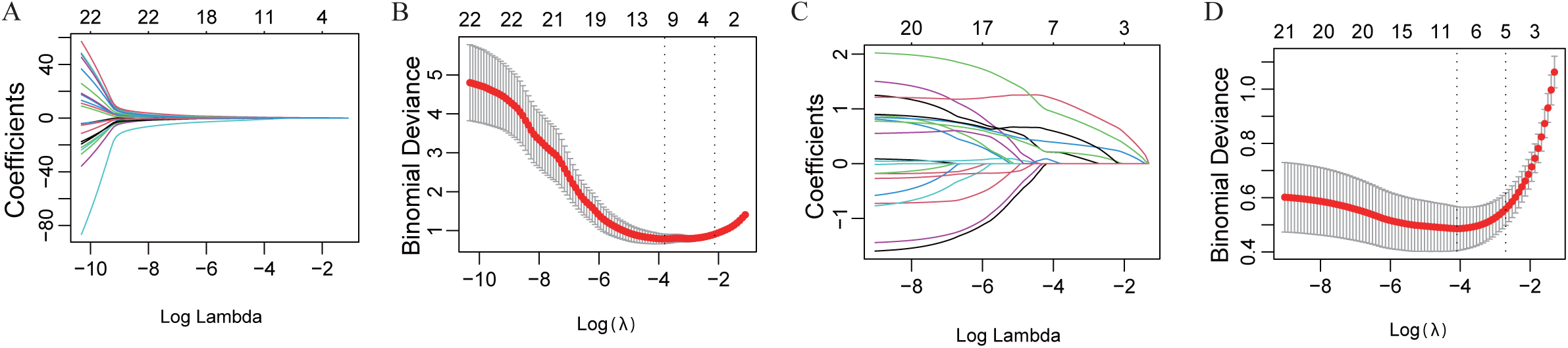
Identification of potential shared diagnostic genes by the LASSO regression model. (A) LASSO coefficient profiles of diagnostic genes in the the AMI database. (B) Tenfold cross-validation to select the optimal tuning parameter log (lambda) in the the AMI database. (C) LASSO coefficient profiles of diagnostic genes in the the PD database. (D) Tenfold cross-validation to select the optimal tuning parameter log (lambda) in the the PD database.

Further investigation showed significant upregulation of FCN1 (Fig. 7A) and LYN (Fig. 7B) in both AMI and PD samples compared with controls. To evaluate their discriminative performance, we performed Receiver Operating Characteristic (ROC) analyses. In the AMI dataset (GSE66360), FCN1 and LYN achieved AUCs of 0.884 and 0.734, respectively (Fig. 7E,F). In the PD dataset (GSE16134), both genes showed strong discrimination with AUCs of 0.879 for FCN1 and 0.882 for LYN (Fig. 7G,H). We also evaluated the two-gene combination. In the AMI dataset, the combined model showed improved discrimination compared with single-gene models (Fig. 7C, p = 3e-04), whereas no additional benefit was observed in the PD dataset (Fig. 7D, p = 0.89556).

**Fig. 7.**
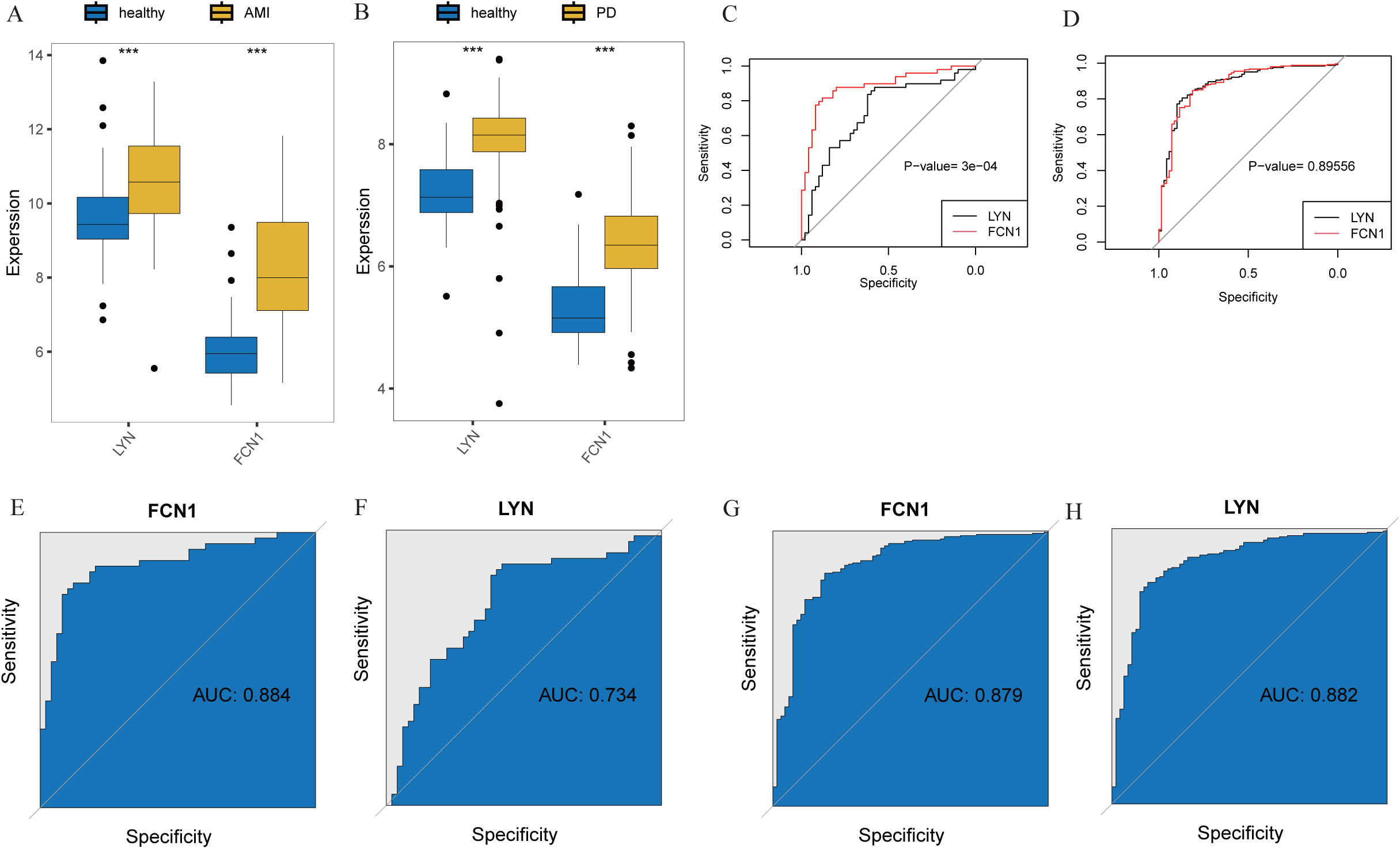
Expression and diagnostic potential of FCN1 and LYN in AMI and PD. (A-B) Box plots showing the expression levels of LYN and FCN1 in healthy controls vs. AMI (A) and healthy controls vs. PD (B). *** indicates p < 0.001. (C-D) Combined ROC curves for LYN and FCN1 in AMI (C) and PD (D) datasets. The p-value indicates the significance of the combined model compared to individual gene models. (E-F) ROC curves showing the diagnostic performance of FCN1 (E) and LYN (F) in the AMI dataset (GSE66360). (G-H) ROC curves demonstrating the diagnostic efficacy of FCN1 (G) and LYN (H) in the PD dataset (GSE16134).

To further evaluate robustness, we assessed FCN1 and LYN in two independent validation datasets (GSE48060 for AMI and GSE10334 for PD), which supported consistent expression patterns and discriminative performance (Supplementary Fig. S1). Collectively, these results suggest that FCN1 and LYN are candidate shared immune markers with potential diagnostic utility for both AMI and PD, warranting further prospective validation.

### Immune infiltration analysis and its correlation with candidate markers

Given the immune-related enrichment results, we explored immune cell patterns using CIBERSORT (Fig. 8). In the PD gingival tissue cohort, plasma cells and neutrophils were increased, whereas resting memory CD4+ T cells were decreased compared with controls (Fig. 8C). In the AMI cohort (GSE66360), immune deconvolution suggested alterations in myeloid- and neutrophil-related fractions (Fig. 8A); however, these results should be interpreted cautiously because GSE66360 profiles CEC-enriched samples rather than bulk whole blood. Correlation analyses indicated that FCN1 was positively associated with monocytes and macrophages M0 signals, and LYN was positively associated with neutrophils and macrophages M0 signals (Fig. 8B,D), supporting a shared myeloid-lineage inflammatory signature.

**Fig. 8.**
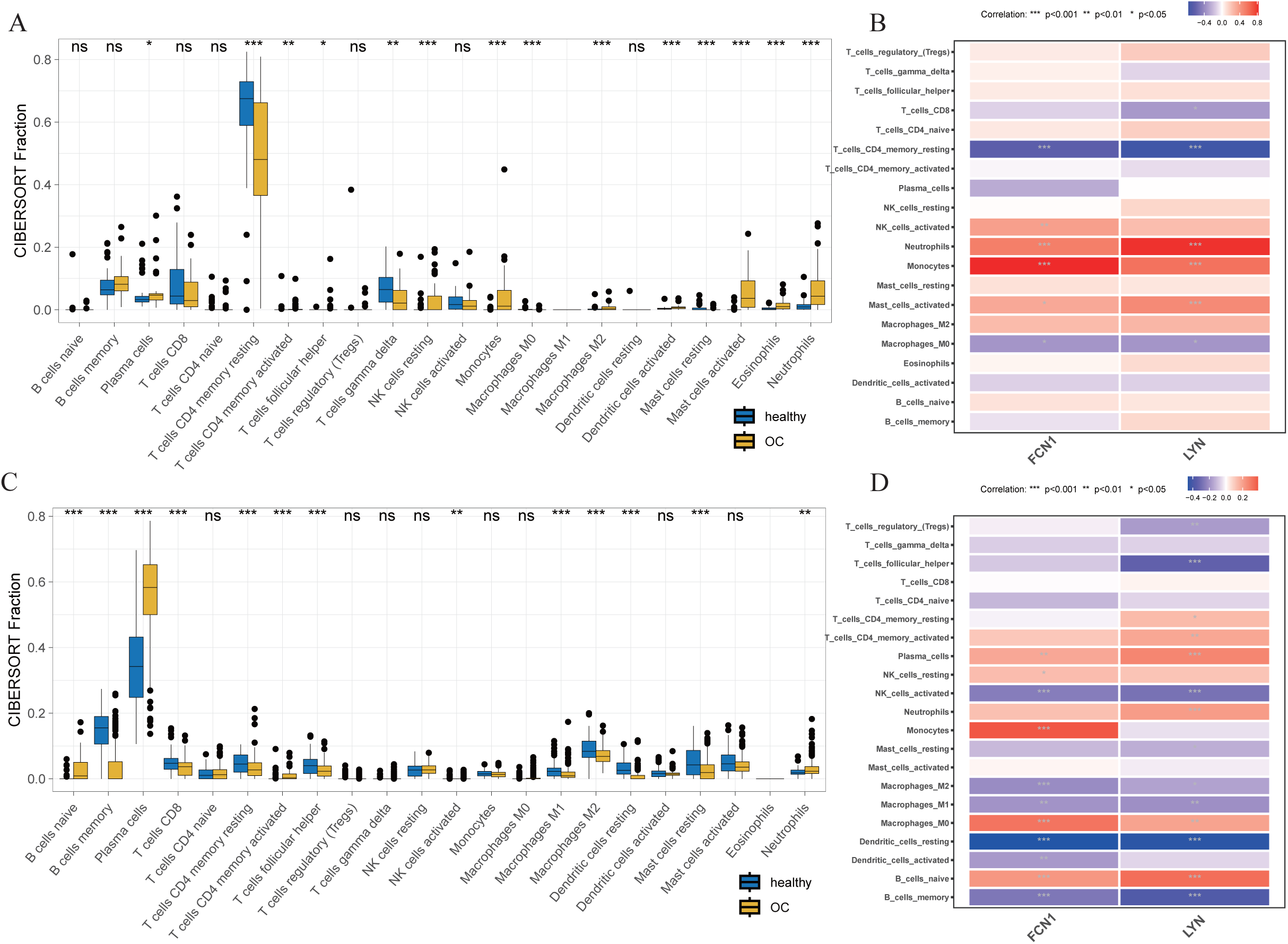
Analysis of immune infiltration associated with AMI and PD. (A) A box plot of the distribution of immune cells in normal samples and AMI samples. (B) The relationship between diagnostic genes and immune cell infiltration in the AMI dataset GSE66360. (C) A box plot of the distribution of immune cells in normal samples and PD samples. (D) The relationship between diagnostic genes and immune cell infiltration in the PD dataset GSE16134.

### Exploring key transcription factors (TFs) and chemicals

To further elucidate the regulatory mechanisms underlying our identified biomarkers, we employed NetworkAnalyst 3.0 (https://www.networkanalyst.ca) to predict the transcription factor (TF)-RNA co-regulatory network. This analysis revealed that STAT3 and GATA2 are targeted by both FCN1 and LYN, suggesting a potential shared regulatory mechanism (Fig. 9A). Additionally, we explored protein-chemical interactions to identify potential drug targets or therapeutic components that could be leveraged for developing interventions addressing both acute AMI and PD simultaneously (Fig. 9B).

**Fig. 9.**
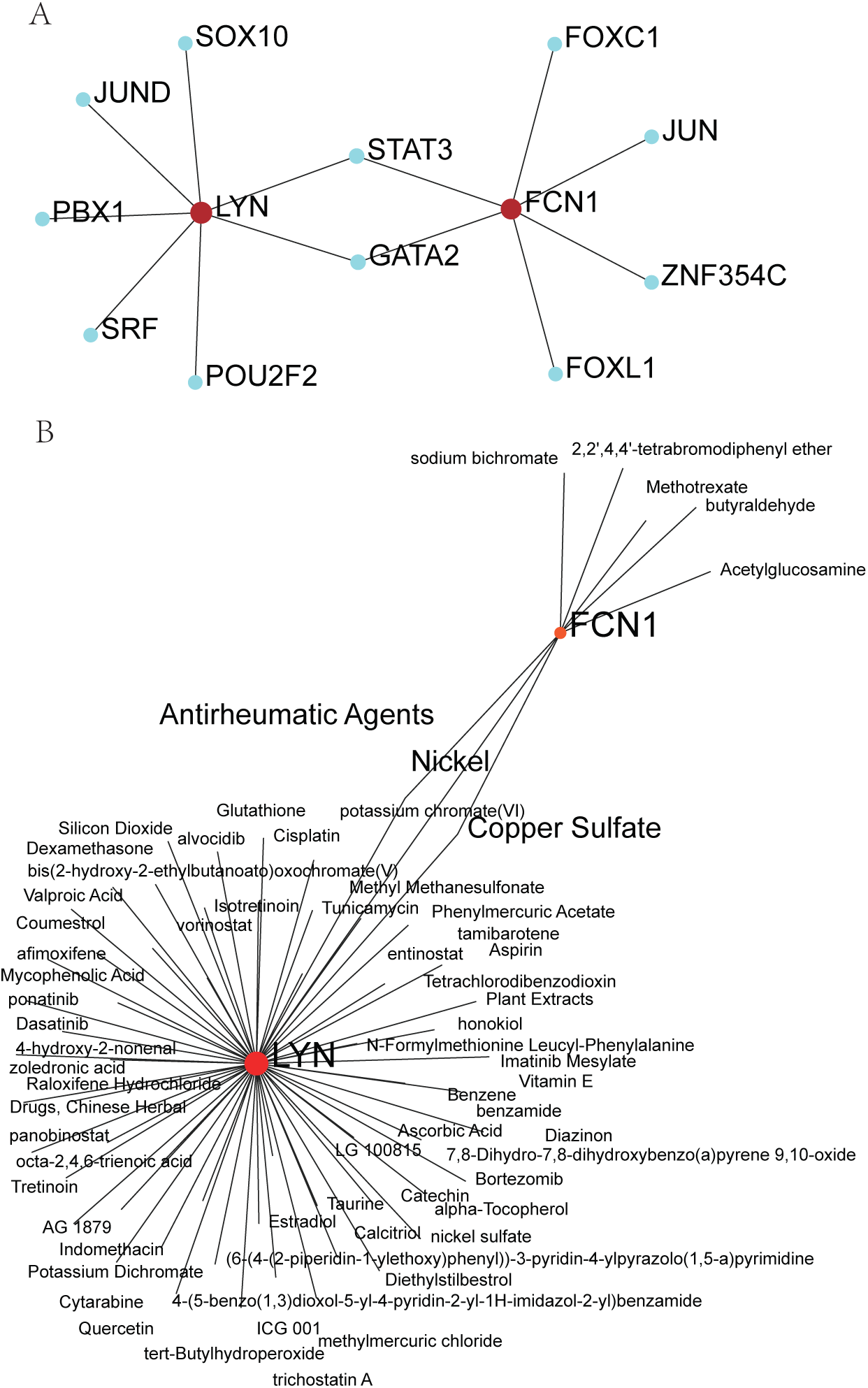
Exploring key transcription factors (TFs) and chemicals. (A) TF-gene interaction network. (B) Protein-chemical interaction network.

### Single-cell analysis of biomarkers locations

To gain deeper insights into the cellular localization of our biomarkers, we performed single-cell analysis on the sample. After quality control, we identified 12 distinct cell clusters (Fig. 10A), which were further categorized into seven major cell populations (Fig. 10B). Subsequent localization analysis of the key diagnostic markers revealed that FCN1 and LYN were predominantly expressed in macrophage:monocyte-derived:M-CSF cells (Fig. 10C-F). This finding provides cellular context for the bulk transcriptomic results and supports a myeloid-associated signature.

**Fig. 10.**
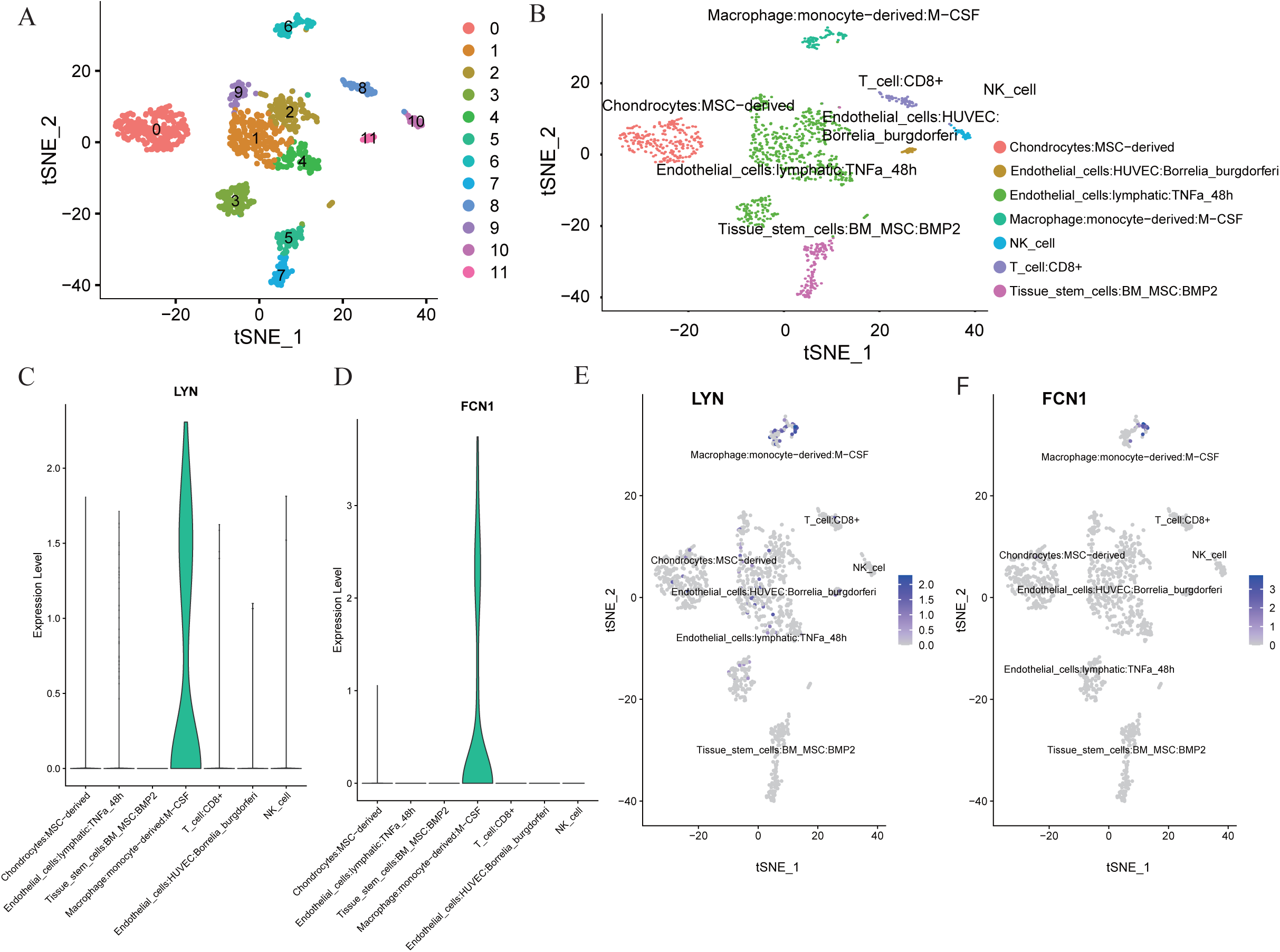
Single-cell analysis. (A) T-SNE visualization of 12 clusters in AMI. (B) T-SNE visualization of cells in AMI. (C,D) Localization analysis of LYN and FCN1. (E,F) The expression levels of LYN and FCN1 in the identified cell types in Myocardium based on the GSE180678 dataset.

## Discussion

AMI remains a leading cause of cardiovascular disease-related mortality and morbidity worldwide, with its precise pathological mechanisms still not fully elucidated ^25^. Interestingly, a growing body of evidence suggests a significant association between AMI and PD, despite their seemingly disparate nature. Epidemiological studies have shown that individuals with PD have an increased risk of developing cardiovascular diseases, including AMI ^26^. A meta-analysis reported that PD was associated with a 1.24-1.35 fold increased risk of coronary heart disease ^27^. The link between oral and cardiovascular health is further supported by the detection of oral pathogens in atherosclerotic plaques ^28,29^. Porphyromonas gingivalis, a key periodontal pathogen, has been found to accelerate atherosclerosis in animal models ^30^. Moreover, treatment of PD has been associated with improvements in endothelial function and reductions in systemic inflammatory markers, suggesting a potential relationship ^31^. Despite these compelling associations, the exact mechanistic links between AMI and PD remain incompletely understood. Our study aims to address this knowledge gap by leveraging bioinformatics approaches to identify shared molecular pathways and potential biomarkers.

Our analysis has revealed molecular and cellular commonalities between AMI and PD, shedding new light on the potential mechanistic links between these seemingly disparate conditions. The identification of 95 shared DEGs, with 91 consistently upregulated in both AMI and PD (Fig. 2C), points to a common inflammatory signature underlying these pathologies. This finding aligns with the growing body of evidence supporting a systemic inflammatory basis for the association between oral and cardiovascular health ^4,6^. The WGCNA further corroborated this shared molecular landscape, revealing 79 overlapping genes in the most positively correlated modules for both diseases (Fig. 3F). This substantial overlap suggests that AMI and PD may activate similar molecular pathways, potentially explaining the epidemiological association observed between these conditions ^7^. Interestingly, the absence of overlap in negatively correlated modules indicates that while AMI and PD may share activation mechanisms, their suppressive or regulatory processes might be distinct. This dichotomy warrants further investigation and could provide insights into disease-specific therapeutic approaches.

Our PPI network analysis identified 22 hub genes (Fig. 4), which were enriched in critical immune and inflammatory processes. The prominence of “leukocyte migration,” “phagocytosis,” and “activation of immune response” in the GO analysis (Fig. 5) underscores the central role of innate immunity in both conditions. This finding is particularly significant in light of recent studies highlighting the importance of neutrophil extracellular traps (NETs) in both cardiovascular disease and periodontitis ^32,33^. NETs, while crucial for pathogen containment, can also contribute to tissue damage and thrombosis when dysregulated, potentially explaining the link between periodontal inflammation and cardiovascular events. The KEGG pathway analysis revealed enrichment in “Lipid and atherosclerosis,” “Staphylococcus aureus infection,” and “Neutrophil extracellular trap formation” pathways (Fig. 5). This result provides a mechanistic bridge between local periodontal inflammation and systemic atherosclerotic processes. The involvement of S. aureus infection pathways is particularly intriguing, as it suggests that oral pathogens or their molecular patterns might trigger similar immune responses in periodontal and cardiovascular tissues. This concept is supported by studies demonstrating the presence of oral bacteria in atherosclerotic plaques ^29,34^.

Our machine learning approach identified FCN1 and LYN as potential shared diagnostic biomarkers for AMI and PD (Fig. 6,7). FCN1, encoding Ficolin-1, is a pattern recognition receptor involved in the lectin pathway of complement activation ^35,36^. Its upregulation in both AMI and PD suggests a potential role in the recognition of damage-associated molecular patterns (DAMPs) or pathogen-associated molecular patterns (PAMPs) in both conditions. LYN, a tyrosine kinase crucial for immune cell signaling, has been implicated in the regulation of inflammatory responses ^37^. The high diagnostic accuracy of these markers (AUCs > 0.7) not only supports their potential as biomarkers but also highlights their importance in the pathogenesis of both diseases.

The immune infiltration analysis revealed increased plasma cells and neutrophils in both AMI and PD (Fig. 8A,C), consistent with the inflammatory nature of these conditions. The elevation of plasma cells is particularly noteworthy, as it suggests a potential role for adaptive immunity in both diseases, possibly through the production of autoantibodies or pathogen-specific antibodies ^38^. The positive correlation of FCN1 and LYN with neutrophils and macrophages (Fig. 8B,D) further supports their role in innate immune responses and provides a cellular context for their action. Our study leveraged a single-cell dataset from an AMI patient to perform annotation analysis, revealing cellular heterogeneity and elucidating underlying mechanisms. We identified seven major cell populations, with FCN1 and LYN expressed to varying degrees predominantly in macrophage:monocyte-derived:M-CSF cells (Fig. 10C-F). This finding aligns with the emerging role of tissue-resident macrophages in both cardiac repair post-AMI and periodontal homeostasis ^39,40^.

We also identified STAT3 and GATA2 as common transcriptional regulators of FCN1 and LYN (Fig. 9A). STAT3, in particular, is known to play crucial roles in both cardiovascular disease and periodontal inflammation ^41,42^. Its involvement suggests that targeting the STAT3 pathway could potentially offer therapeutic benefits for both conditions simultaneously. The protein-chemical interaction analysis (Fig. 9B) opens up exciting possibilities for drug repurposing or the development of novel therapeutic strategies that could address both AMI and PD. This approach could lead to more holistic treatment paradigms that consider the systemic nature of these seemingly localized diseases.

In conclusion, our multi-faceted analysis identifies FCN1 and LYN as key genes and reveals a shared inflammatory and immune-mediated pathogenic signature between AMI and PD. This signature is centered on neutrophil and macrophage activation, complement pathways, and specific transcriptional regulators. The identified biomarkers and pathways not only provide potential diagnostic tools but also suggest novel therapeutic targets that could address both conditions simultaneously. These findings underscore the importance of considering oral health in cardiovascular disease prevention and vice versa, potentially leading to new integrated approaches in patient care and public health strategies.

### Limitations

Our study offers several strengths. However, several limitations must be acknowledged. The retrospective nature of our data, sourced from public databases, may introduce biases; future studies should aim for prospective data collection to enhance predictive power. The limited sample size, particularly for single-cell analysis, may not fully represent the broader population, necessitating validation in larger, more diverse cohorts. Additionally, while our bioinformatics approach provides valuable insights, experimental validation is crucial to confirm the functional roles of identified genes and pathways. Furthermore, our models, while informative, are not infallible and may not account for all relevant factors such as disease severity, comorbidities, and population-specific variations. Despite these limitations, our study provides a comprehensive molecular perspective on the AMI-PD relationship, laying a foundation for future investigations and potential therapeutic interventions.

## Acknowledgement

We would like to express our appreciations to the participants and their families.

## Availability of data and materials

This research was conducted using the GEO database. Data are publicly available from www.ncbi.nlm.nih.gov/geo. We would like to express our appreciations to the participants and their families.

## Ethics approval and consent to participate

Not applicable. **Consent to participate** Not applicable.

## Consent for publication

Not applicable.

## Authors’ contributions

K.Z.: Data curation, Software, Visualization, Methodology, Original draft preparation. YQ.W.: Conceptualization, Supervision, Funding acquisition, Writing and Editing.

---

**Supplementary Fig S1** Expression pattern validation and diagnostic value in validation cohorts. (A) Box plots showing the expression levels of LYN and FCN1 in healthy controls vs. AMI patients in the validation cohort GSE48060. (B) Box plots depicting the expression levels of LYN and FCN1 in healthy controls vs. PD patients in the validation cohort GSE10334. (C) ROC curves demonstrating the diagnostic performance of FCN1 and LYN in the AMI validation dataset GSE48060. AUC values are provided for each gene. (D) ROC curves illustrating the diagnostic efficacy of FCN1 and LYN in the PD validation dataset GSE10334. AUC values are shown for each gene.

## References

1. Virani SS, Alonso A, Aparicio HJ, et al. Heart Disease and Stroke Statistics-2021 Update: A Report From the American Heart Association. Circulation 2021; 143(8): e254–e743.

2. Wang K, Jin Y, Wang M, et al. Global cardiovascular diseases burden attributable to high sodium intake from 1990 to 2019. 2023; 25(9): 868–79.

3. Yusuf S, Joseph P, Rangarajan S, et al. Modifiable risk factors, cardiovascular disease, and mortality in 1558722 individuals from 21 high-income, middle-income, and low-income countries (PURE): a prospective cohort study. Lancet 2020; 395(10226): 795–808.

4. Tonetti MS, Van Dyke TE. Periodontitis and atherosclerotic cardiovascular disease: consensus report of the Joint EFP/AAP Workshop on Periodontitis and Systemic Diseases. Journal of periodontology 2013; 84(4 Suppl): S24–9.

5. Alvarenga MOP, Frazão DR, de Matos IG, et al. Is There Any Association Between Neurodegenerative Diseases and Periodontitis? A Systematic Review. Frontiers in aging neuroscience 2021; 13: 651437.

6. Hajishengallis G. Periodontitis: from microbial immune subversion to systemic inflammation. Nat Rev Immunol 2015; 15(1): 30–44.

7. Sanz M, Marco Del Castillo A, Jepsen S, et al. Periodontitis and cardiovascular diseases: Consensus report. Journal of clinical periodontology 2020; 47(3): 268–88.

8. Dietrich T, Sharma P, Walter C, Weston P, Beck J. The epidemiological evidence behind the association between periodontitis and incident atherosclerotic cardiovascular disease. Journal of clinical periodontology 2013; 40 Suppl 14: S70–84.

9. Beukers NG, van der Heijden GJ, van Wijk AJ, Loos BG. Periodontitis is an independent risk indicator for atherosclerotic cardiovascular diseases among 60 174 participants in a large dental school in the Netherlands. J Epidemiol Community Health 2017; 71(1): 37–42.

10. Leng Y, Hu Q, Ling Q, et al. Periodontal disease is associated with the risk of cardiovascular disease independent of sex: A meta-analysis. Front Cardiovasc Med 2023; 10: 1114927.

11. Rydén L, Buhlin K, Ekstrand E, et al. Periodontitis Increases the Risk of a First Myocardial Infarction: A Report From the PAROKRANK Study. Circulation 2016; 133(6): 576–83.

12. Libby P, Loscalzo J, Ridker PM, et al. Inflammation, Immunity, and Infection in Atherothrombosis: JACC Review Topic of the Week. J Am Coll Cardiol 2018; 72(17): 2071–81.

13. Winning L, Linden GJ. Periodontitis and Systemic Disease: Association or Causality? Curr Oral Health Rep 2017; 4(1): 1–7.

14. Barrett T, Wilhite SE, Ledoux P, et al. NCBI GEO: archive for functional genomics data sets--update. Nucleic acids research 2013; 41(Database issue): D991–5.

15. Ritchie ME, Phipson B, Wu D, et al. limma powers differential expression analyses for RNA-sequencing and microarray studies. Nucleic acids research 2015; 43(7): e47.

16. Ito K, Murphy D. Application of ggplot2 to Pharmacometric Graphics. CPT: pharmacometrics & systems pharmacology 2013; 2(10): e79.

17. Chen H, Boutros PC. VennDiagram: a package for the generation of highly-customizable Venn and Euler diagrams in R. BMC bioinformatics 2011; 12: 35.

18. Stuart T, Butler A, Hoffman P, et al. Comprehensive Integration of Single-Cell Data. Cell 2019; 177(7): 1888–902.e21.

19. Langfelder P, Horvath S. WGCNA: an R package for weighted correlation network analysis. BMC bioinformatics 2008; 9: 559.

20. Shannon P, Markiel A, Ozier O, et al. Cytoscape: a software environment for integrated models of biomolecular interaction networks. Genome research 2003; 13(11): 2498–504.

21. Gene Ontology Consortium: going forward. Nucleic acids research 2015; 43(Database issue): D1049–56.

22. Xie Y, Shi H, Han B. Bioinformatic analysis of underlying mechanisms of Kawasaki disease via Weighted Gene Correlation Network Analysis (WGCNA) and the Least Absolute Shrinkage and Selection Operator method (LASSO) regression model. BMC pediatrics 2023; 23(1): 90.

23. Newman AM, Liu CL, Green MR, et al. Robust enumeration of cell subsets from tissue expression profiles. Nature methods 2015; 12(5): 453–7.

24. Zhou G, Soufan O, Ewald J, Hancock REW, Basu N, Xia J. NetworkAnalyst 3.0: a visual analytics platform for comprehensive gene expression profiling and meta-analysis. Nucleic acids research 2019; 47(W1): W234–w41.

25. Frangogiannis NG. Pathophysiology of Myocardial Infarction. Comprehensive Physiology 2015; 5(4): 1841–75.

26. Lockhart PB, Bolger AF, Papapanou PN, et al. Periodontal disease and atherosclerotic vascular disease: does the evidence support an independent association?: a scientific statement from the American Heart Association. Circulation 2012; 125(20): 2520–44.

27. Humphrey LL, Fu R, Buckley DI, Freeman M, Helfand M. Periodontal disease and coronary heart disease incidence: a systematic review and meta-analysis. J Gen Intern Med 2008; 23(12): 2079–86.

28. Kebschull M, Demmer RT, Papapanou PN. “Gum bug, leave my heart alone!”--epidemiologic and mechanistic evidence linking periodontal infections and atherosclerosis. Journal of dental research 2010; 89(9): 879–902.

29. Chhibber-Goel J, Singhal V, Bhowmik D, et al. Linkages between oral commensal bacteria and atherosclerotic plaques in coronary artery disease patients. NPJ Biofilms Microbiomes 2016; 2: 7.

30. Hayashi C, Viereck J, Hua N, et al. Porphyromonas gingivalis accelerates inflammatory atherosclerosis in the innominate artery of ApoE deficient mice. Atherosclerosis 2011; 215(1): 52–9.

31. Tonetti MS, D’Aiuto F, Nibali L, et al. Treatment of periodontitis and endothelial function. N Engl J Med 2007; 356(9): 911–20.

32. Jorch SK, Kubes P. An emerging role for neutrophil extracellular traps in noninfectious disease. Nat Med 2017; 23(3): 279–87.

33. Delbosc S, Alsac JM, Journe C, et al. Porphyromonas gingivalis participates in pathogenesis of human abdominal aortic aneurysm by neutrophil activation. Proof of concept in rats. PLoS One 2011; 6(4): e18679.

34. Koren O, Spor A, Felin J, et al. Human oral, gut, and plaque microbiota in patients with atherosclerosis. Proc Natl Acad Sci U S A 2011; 108 Suppl 1(Suppl 1): 4592–8.

35. Endo Y, Matsushita M, Fujita T. New insights into the role of ficolins in the lectin pathway of innate immunity. Int Rev Cell Mol Biol 2015; 316: 49–110.

36. Endo Y, Matsushita M, Fujita T. The role of ficolins in the lectin pathway of innate immunity. Int J Biochem Cell Biol 2011; 43(5): 705–12.

37. Scapini P, Pereira S, Zhang H, Lowell CA. Multiple roles of Lyn kinase in myeloid cell signaling and function. Immunol Rev 2009; 228(1): 23–40.

38. Tsiantoulas D, Diehl CJ, Witztum JL, Binder CJ. B cells and humoral immunity in atherosclerosis. Circ Res 2014; 114(11): 1743–56.

39. Epelman S, Liu PP, Mann DL. Role of innate and adaptive immune mechanisms in cardiac injury and repair. Nat Rev Immunol 2015; 15(2): 117–29.

40. Dutzan N, Konkel JE, Greenwell-Wild T, Moutsopoulos NM. Characterization of the human immune cell network at the gingival barrier. Mucosal Immunol 2016; 9(5): 1163–72.

41. Hirano T, Ishihara K, Hibi M. Roles of STAT3 in mediating the cell growth, differentiation and survival signals relayed through the IL-6 family of cytokine receptors. Oncogene 2000; 19(21): 2548–56.

42. Mao S, Park Y, Hasegawa Y, et al. Intrinsic apoptotic pathways of gingival epithelial cells modulated by Porphyromonas gingivalis. Cell Microbiol 2007; 9(8): 1997–2007.

