## Supplementary figures and images for "Shared immune signatures in acute myocardial infarction and periodontitis: the role of FCN1 and LYN as potential biomarkers"

### Supplementary Figure 1

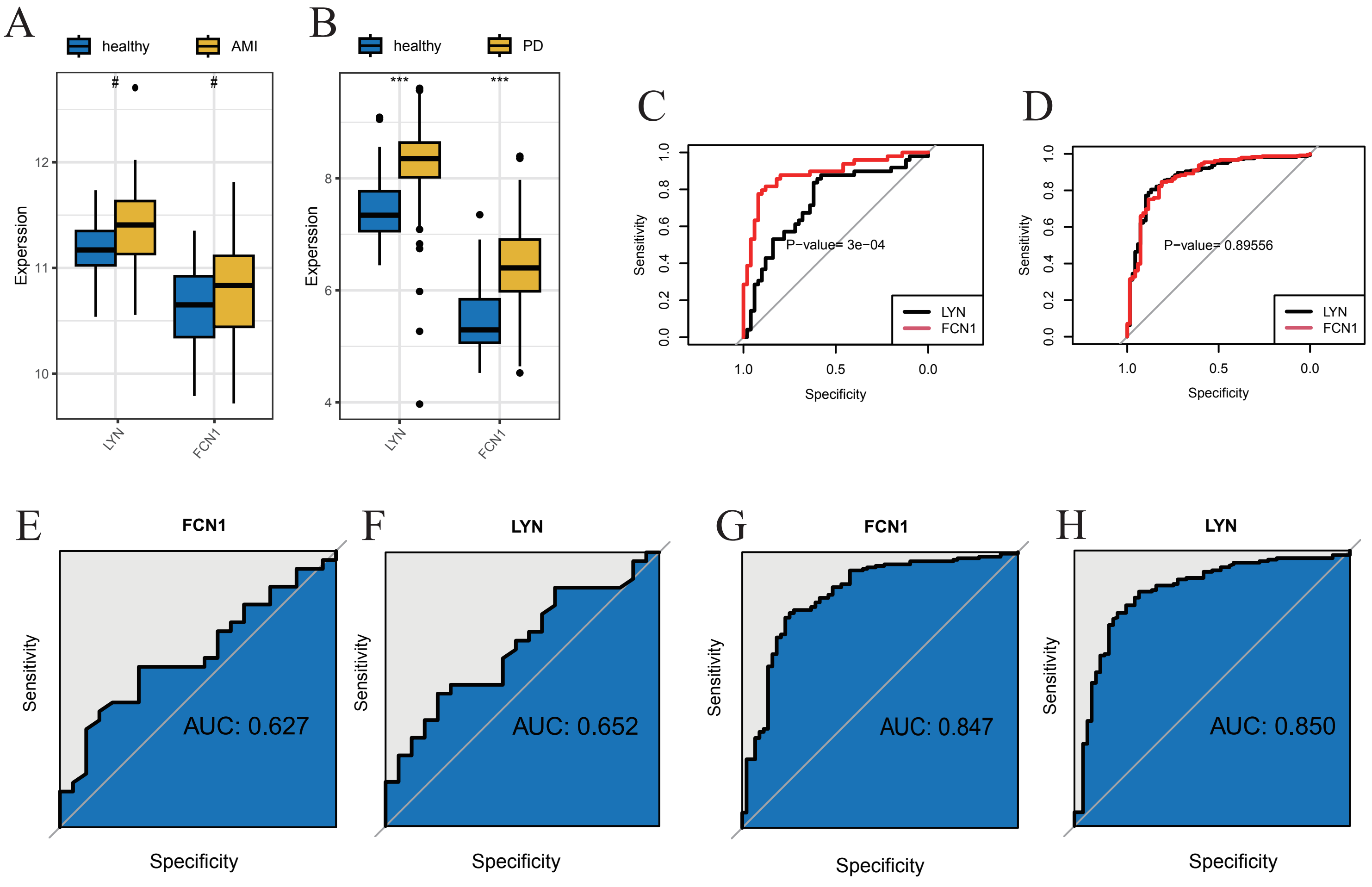
